# *Escherichia coli* populations adapt to complex, unpredictable fluctuations without any trade-offs across environments

**DOI:** 10.1101/045369

**Authors:** Shraddha Karve, Devika Bhave, Dhanashri Nevgi, Sutirth Dey

## Abstract

In nature, organisms are simultaneously exposed to multiple stresses (i.e. complex environments) that often fluctuate unpredictably. While both these factors have been studied in isolation, the interaction of the two remains poorly explored. To address this issue, we selected laboratory populations of *Escherichia coli* under complex (i.e. stressful combinations of pH, H_2_O_2_ and NaCl) unpredictably fluctuating environments for ~900 generations. We compared the growth rates and the corresponding trade-off patterns of these populations to those that were selected under constant values of the component stresses (i.e. pH, H_2_O_2_ and NaCl) for the same duration. The fluctuation-selected populations had greater mean growth rate and lower variation for growth rate over all the selection environments experienced. However, while the populations selected under constant stresses experienced severe tradeoffs in many of the environments other than those in which they were selected, the fluctuation-selected populations could by-pass the across-environment trade-offs completely. Interestingly, trade-offs were found between growth rates and carrying capacities. The results suggest that complexity and fluctuations can strongly affect the underlying trade-off structure in evolving populations.

## INTRODUCTION

Although there is considerable understanding on how populations evolve under a single selection environment (reviewed in Scheiner 2002; Prasad and Joshi 2003; Lenski 2004), most natural populations face temporally fluctuating complex environments (i.e. simultaneous exposure to multiple stresses). Over the last few decades, a large number of theoretical (Levins 1968; Whitlock 1996; Lande 2008) and empirical (Leroi et al. 1994; Reboud and Bell 1997; Hallsson and Björklund 2012; Alto et al. 2013; Ketola et al. 2013; Condon et al. 2014; Karve et al. 2015; Karve et al. 2016) studies have investigated the fitness outcomes of different kinds of environmental complexities and fluctuations. For example, microbial populations facing predictable environmental fluctuations show improved fitness over the entire range of fluctuating environments (Leroi et al. 1994; Hughes et al. 2007; Coffey and Vignuzzi 2011). On the other hand, unpredictable fluctuations sometimes improve fitness of the microbes in all environments (Turner and Elena 2000; Ketola et al. 2013) or improve in some of the environments without any change in others (Hughes et al. 2007). Similarly, complex resource environments also result in fitness improvements, albeit to different degrees, for all the resources experienced during selection (Barrett et al. 2005; Cooper and Lenski 2010). Unlike mean fitness though, variation for fitness shows opposing trends for fluctuating and complex environments. Populations selected under fluctuations often show reduced variation for fitness over all the selection environments (Kassen 2014 Table 4.2) while complex resource environments tend to increase the variation for fitness across resources (Barrett et al. 2005). However the evolutionary outcomes of simultaneously facing environmental fluctuations and complexity remain relatively unexplored in terms of mean fitness as well as variation for fitness. This lack of understanding also extends to the underlying constraints experienced by evolving populations, particularly in the context of the patterns of trade-offs.

Trade-offs, i.e. a gain of fitness in some environment(s), accompanied by loss of fitness in other environment(s), can be observed between two different life history traits in a given environment or between two different environments for a given life history trait (Agrawal et al. 2010). The latter is thought to be more relevant in microbial systems where there are very few “life-history traits” and most fitness measurements involve composite traits like growth rate or competitive ability. Interestingly, in microbial experimental evolution studies, tradeoffs are mostly observed in novel environments, i.e. environments not experienced during selection (Bennett et al. 1992; Reboud and Bell 1997; Hughes et al. 2007; Jasmin and Kassen 2007) and rarely across selection environments. This suggests that accumulation of conditionally neutral mutations, and not antagonistic pleiotropic interactions, are chiefly responsible for trade-offs in microbes (Kassen 2014 Figure 3.6). However, it is difficult to predict how such mutations would accumulate in the face of environmental complexity and unpredictable fluctuations, and not surprisingly, little is known about the patterns of trade-off in microbial populations exposed to such conditions.

Here we study the evolutionary implications of the interaction of complexity and temporal fluctuations in environments, Replicate laboratory populations of *Escherichia coli* were exposed to complex (i.e. stressful combinations of pH, H_2_O_2_ and NaCl), unpredictably fluctuating environments. Parallely, we also selected replicate bacterial populations under constant exposure to each of the selection environments. After ~900 generations of selection, we found that populations facing complex, unpredictable environments increased their mean fitness (measured as population growth rates) and minimized the variation for fitness when estimated over all the selection environments. Moreover, selection under complex, fluctuating environments did not lead to trade-offs across different selection environments, although some trade-offs were observed between growth rate and carrying capacity in some of the environments. Our results show that populations facing unpredictably fluctuating complex environments can show higher mean fitness and significantly lower fitness variation without undergoing any trade-offs across selection environments.

## METHODS

### Selection experiment

We used Kanamycin resistant *Escherichia coli* strain K12 (see Supplementary online material S1 for details) for this study. A single colony grown on Nutrient agar with Kanamycin (see SOM for composition) was inoculated in 2 ml of Nutrient broth with Kanamycin (NB^Kan^) (see Supplementary online material S2 for composition) and allowed to grow for 24 h at 37^0^C,150 rpm in 24 welled plate. 4 μl of this suspension was used to initiate each of 120 replicate populations.

120 replicate populations were equally divided into five treatment regimes and one control regime, such that there were 20 replicate populations per regime. Control populations were subcultured in NB^Kan^ for the entire duration of the selection. Four out of five selection regimes were constant environments with salt or pH9 or pH4.5 or hydrogen peroxide (H_2_O_2_) in NB^Kan^. The remaining selection environment was complex and stochastically fluctuating (henceforth termed as F) (see Supplementary online material S3 for details of all selection regimes). Detailed design of this fluctuating selection regime has been mentioned elsewhere (Karve et al. 2015).

24 welled plates with 2 ml of appropriate growth medium and 4 μl of inoculum volume for each well were used throughout the selection and assay experiments. The growth conditions were maintained at 37^0^C, 150 rpm. All the populations were sub-cultured every 24 h. Extinctions were identified visually as a lack of turbidity and revived using 20 μl of the previous day’s culture stored at 4^0^C. The selection lasted for 100 days i.e. ~900 generations, computed using the standard expression for calculation of generation time (Bennett and Lenski 1997). On the 100^th^ day, the populations were stored as glycerol stocks at −80^0^C for future assays.

### Fitness assay in selection environments

After 100 days of selection, all the populations were assayed for fitness in every selection environment, except the fluctuating environment. Selection environments comprised of NB^Kan^ with salt or pH9 or pH4.5 or hydrogen peroxide or control (see Supplementary online material S3 for details).

For the comparison with the ancestor, we revived the ancestral population of Kanamycin resistant *Escherichia coli* strain K12 in NB^Kan^ for the duration of 18 hr. 20 replicate wells were inoculated with this revived culture for every selection environment. This resulted in the same number of replicates of ancestral culture for every assay environment as that of the selected populations.

Following previous studies, maximum growth rate during 24 h of growth was used as a fitness measure (Ketola et al. 2013; Karve et al. 2015). For the growth rate measurement, 4 μl of relevant glycerol stocks were revived in 2 ml of in NB^Kan^. After 18 h of growth, these revived cultures were inoculated in the appropriate assay environment. OD_600_ was measured every 2 h on a plate reader (Synergy HT BioTek, Winooski, VT, USA) for the duration of 24 h. We used a QBASIC (v 4.5) script to determine the maximum growth rate of the bacterial populations. The program fits a straight line on overlapping moving windows of three points on the time series of OD_600_ values. The maximum slope obtained by this method was taken as the maximum growth rate for that population.

We used the same 24 h growth trajectories, to estimate the maximum density reached (henceforth carrying capacity), which was used as an alternate proxy for fitness. Both the fitness estimates, i.e. maximum growth rate and maximum density reached, were analyzed as mentioned below.

### Replicates and statistical analysis

For every population (6 selection regimes × 20 replicates + 20 replicates of the ancestral population) growth assays were performed twice in five environments (pH4.5, pH9, H_2_O_2_, salt, NB). This resulted in a total of 1400 growth measurements (140 populations × 5 assay environments × 2 measurements).

### Overall mean fitness

1200 fitness estimates (6 selection regimes × 5 assay environments × 20 replicates × 2 measurements, excluding the ancestor) were analysed using a 3-way mixed model ANOVA. Selection (six levels) and assay environment (5 levels) were fixed factors crossed with each other. Replicate (twenty levels) was taken as a random factor nested in selection. To determine whether F populations differed significantly from the other selection lines, we performed Dunnett’s post hoc test (Zar 1999).

### Variation for fitness

We estimated coefficients of variation (CV) as a measure of variation in fitness. Two fitness estimates in a given assay environment were averaged and CV was calculated for every replicate population over this average fitness in five assay environments. Every selection regime thus yielded 20 CV estimates (one for each replicate population). These were then analyzed using a one way ANOVA with selection (six levels) as a fixed factor. To determine whether F populations differ significantly from other selection lines, we performed Dunnett’s post hoc test (Zar 1999).

### Trade off

The mean fitness of the ancestor for every selection environment was computed as the average of 40 measurements (20 replicates × 2 measurements). This difference was then subtracted from the mean fitness for every population in the same environment. This resulted in 20 difference measurements for every selection regime in every selection environment.

We then performed 30 different t tests (6 selection lines × 5 assay environments) for every set of difference computed from ancestor. Family-wise error rates were controlled through sequential Holm-Šidàk correction of the *p*-values (Abdi 2010).

To estimate the biological significance of the difference in fitness of F populations compared to other selected populations, we computed Cohen’s *d* statistics (Cohen 1988) as a measure of effect size. It was interpreted as small, medium and large for 0.2 < d < 0.5, 0.5 < d < 0.8 and d > 0.8, respectively.

All the ANOVAs were performed on STATISTICA v7.0 (Statsoft Inc.). Cohen’s *d* statistics were estimated using the freeware Effect size generator v2.3.0 (Devilly 2004).

## RESULTS

### Fluctuating environments select for higher overall mean fitness and lower variation for fitness

When pooled over all the selection environments, effect of selection was highly significant (*F*_16, 700_ = 12.07, *p* < 0.0001). The effect of assay environment was also highly significant (*F*_4, 700_ = 247.04, *p* < 0.001) and there was a significant interaction of selection with assay environment (*F*_24, 600_ = 31.73*,p* < 0.001). The growth rates of the selected populations vis-avis the ancestor were compared using the Dunnett’s test. F populations showed the highest overall mean fitness which was significantly greater that the mean fitness of ancestor with medium effect size (Table 1, Fig 1A). None of the other selected populations differed significantly from the ancestor.

**Table 1.**
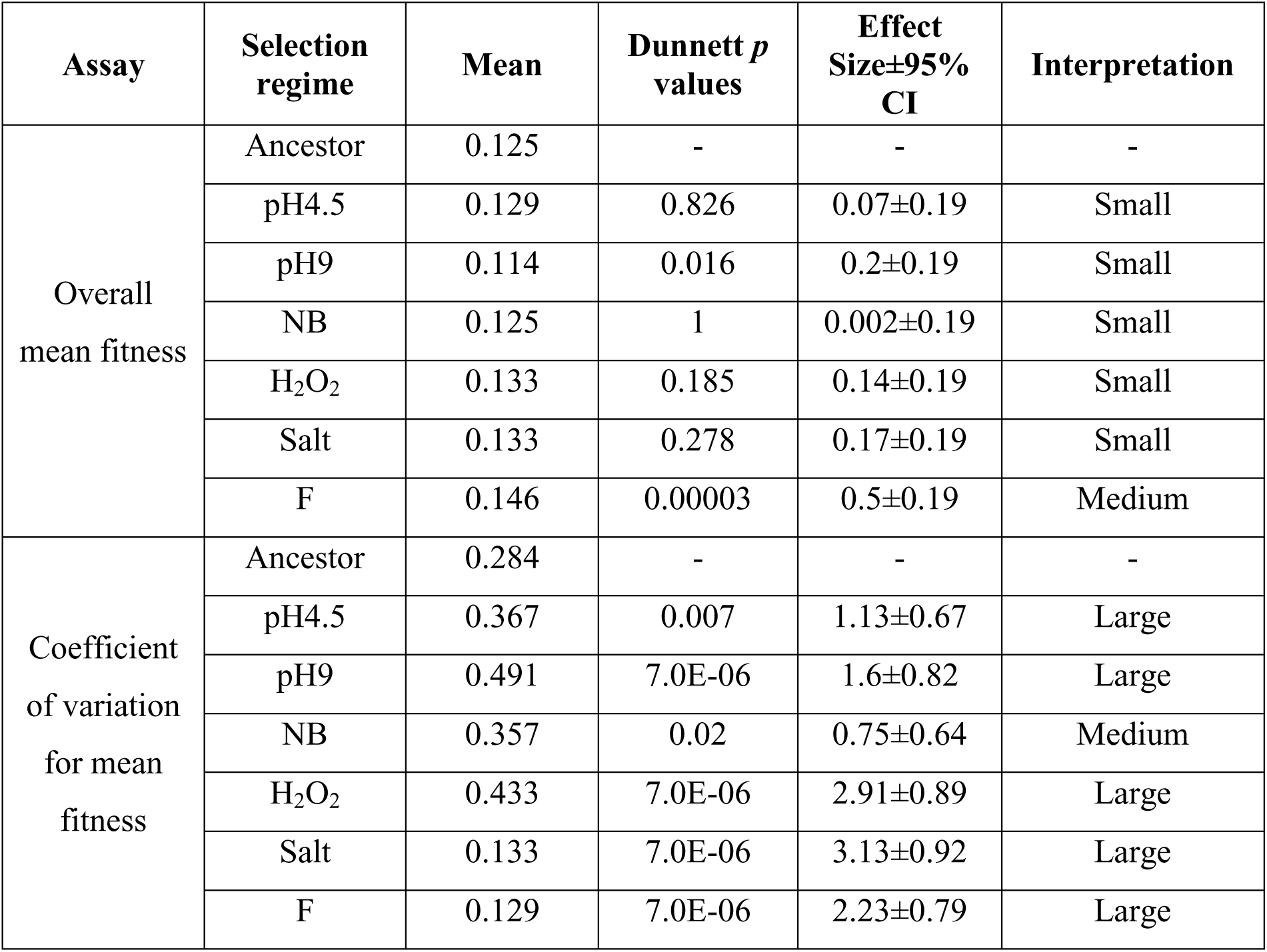
Summary of the main effect of selection in the ANOVAs for individual selection regimes. Dunnett’s post hoc test was conducted with ancestor as a control group. Dunnett’s test *p* values and effect size are thus in comparison with ancestor.

**Figure 1:**
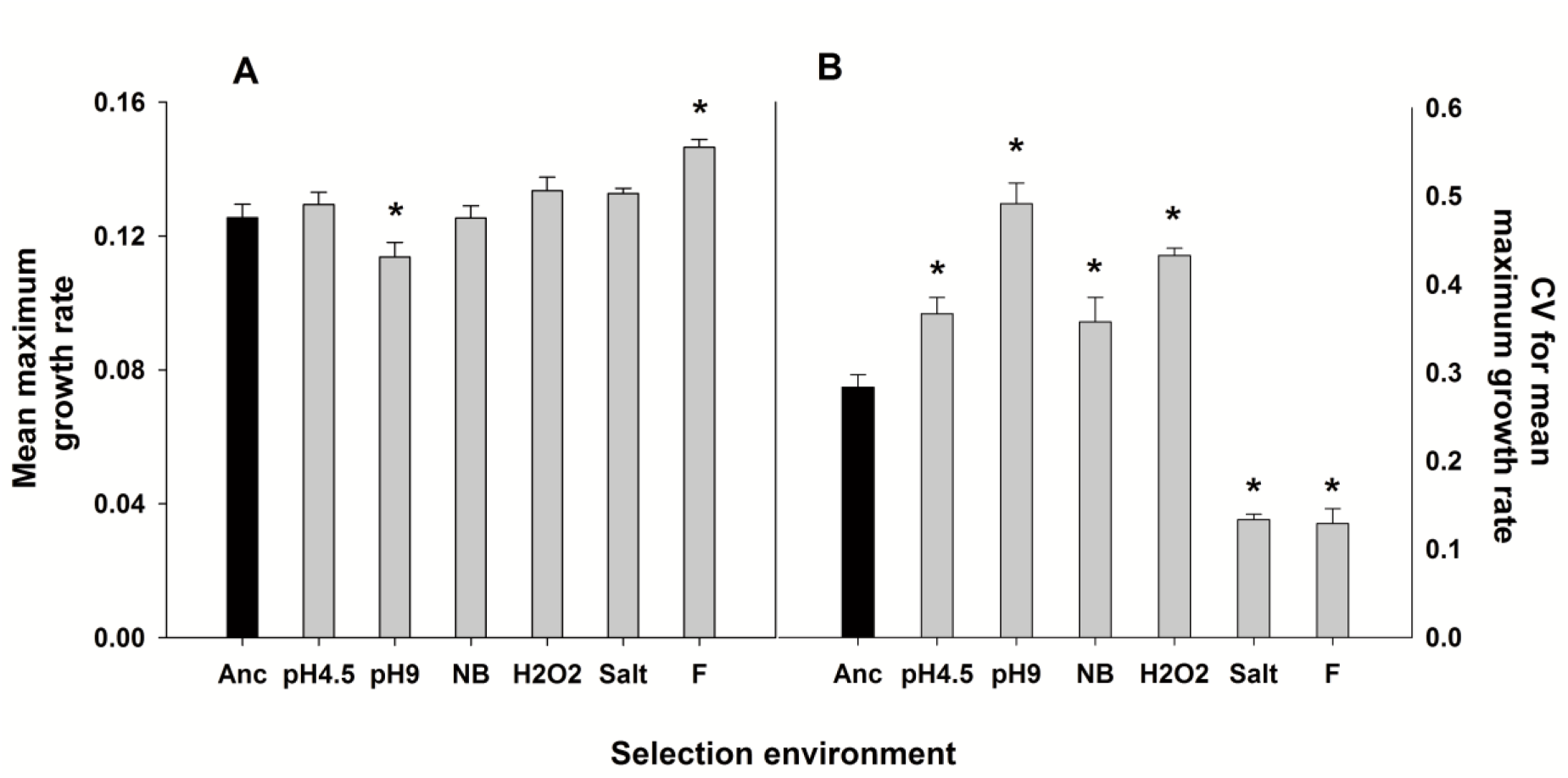
Mean fitness and CV for mean fitness for all the selection regimes. Fitness estimated as maximum slope of the growth trajectory over 24 hours. Overall mean fitness was computed for every selection regime over all assay environments. Coefficient of variation was computed for every selection regime over all the assay environments. Error bars represent SEM. * denotes significantly (*p* < 0.05, Dunnett’s post hoc test) different mean fitness or mean CV for fitness, from ancestor. The F populations had the highest mean fitness and the lowest CV for fitness across all the assay regimes.

Selection also had a significant effect on the variation of fitness across all environments (*F*_6, 133_ = 61.65, *p* < 0.0001). The coefficient of variation of the F populations was the lowest among all the treatments and it was significantly lower than the ancestor with a large effect size (Table 1, Fig 1B). The populations selected in salt also had significantly lower variation for fitness (compared to the ancestor) with large effect size (Table 1, Fig 1B). All the other populations had significantly greater variation in fitness than the ancestor across all environments.

To summarize, these results suggest that selection in fluctuating environments leads to a modest increase in overall mean fitness along with lower variation for fitness. Populations selected in salt also minimize the variation effectively but show loss of fitness as compared to the ancestor.

### Fluctuating environments minimize trade-offs in growth rates

F populations did not show any significant loss in fitness compared to the ancestor, in any of the selection environments (Fig 2). In contrast, populations selected in constant environments always showed a significant loss of fitness in at least one of the selection environments (Table S5 (A), Fig 2). All the selected populations showed significant increase in fitness in hydrogen peroxide and all of them, except F populations, show loss of fitness in salt. Interestingly, even the populations selected in salt showed reduction in fitness when assayed in salt, albeit to a lesser degree (see discussion).

**Figure 2:**
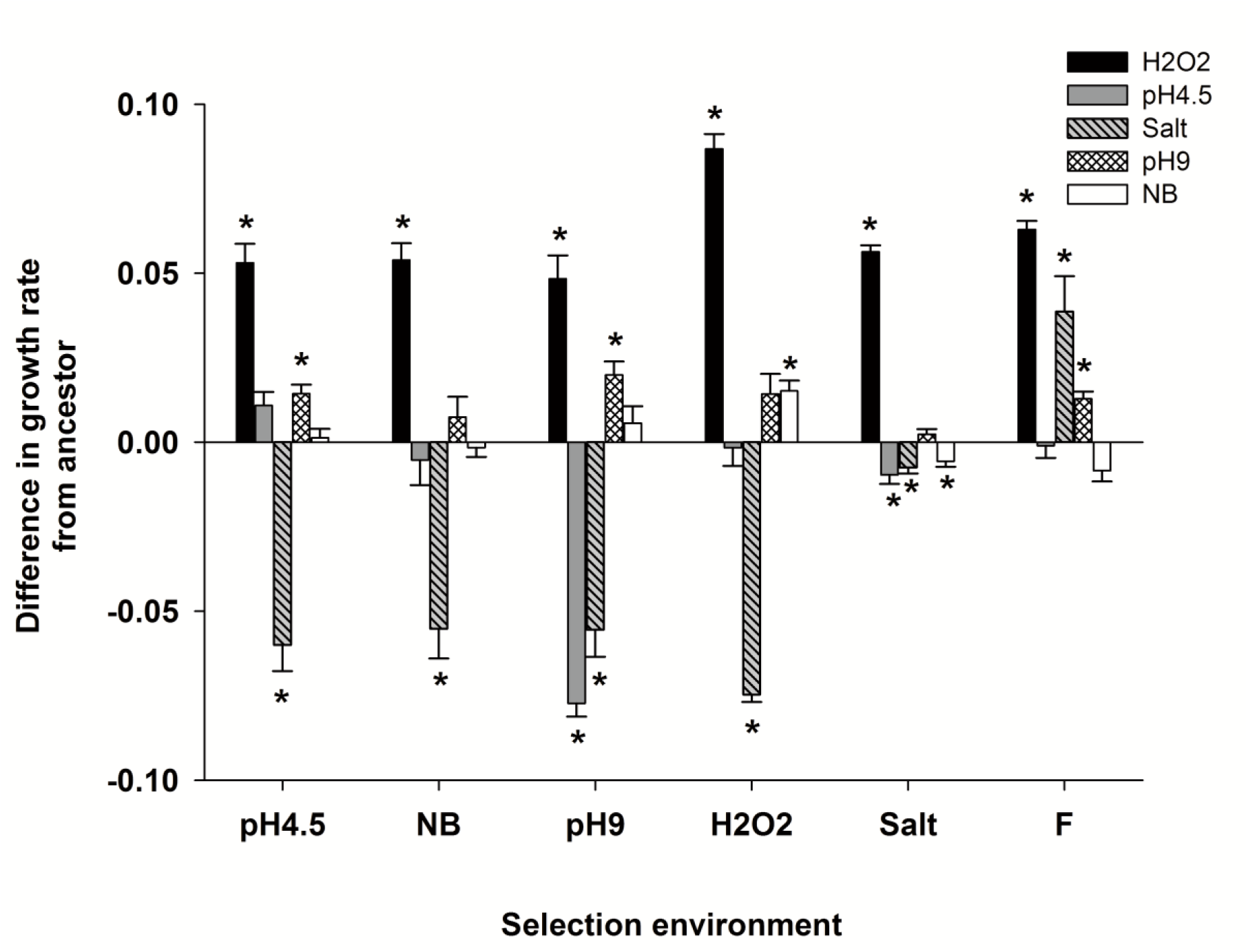
Difference (±SE) in maximum growth rate from ancestor for all the selection regimes. Difference in mean fitness (estimated as maximum growth rate) from ancestor was computed for every selection regime in every selection environment. Negative values indicate loss of fitness while positive values indicate gain of fitness, as compared to the ancestor. Every selection regime, except F, shows loss of fitness in at least one of the environments. * denotes *p* < 0.05 after individual ANOVAs followed by Holm-Šidák correction.

Taken together, these results show that fluctuating environments can minimize trade-offs.

### Trade-offs can be seen across two different measures of fitness

When we analyzed the mean and variance in fitness estimated as carrying capacity, the patterns in overall mean and variance were similar to those obtained with maximum growth rate (Supplementary online material S4), i.e. F populations showed the highest mean fitness with lowest variation for fitness. Compared to the ancestors, F populations improved fitness in every selection environment with an exception of pH4.5 (Fig 3). Interestingly, populations selected in salt showed improved carrying capacity in salt as well as NB even though there was a significant reduction in maximum growth rate in these two assay environments. These results suggest that apart from trade-offs across environments, microbial populations can also exhibit trade-offs across different measures of fitness in a given environment.

**Figure 3:**
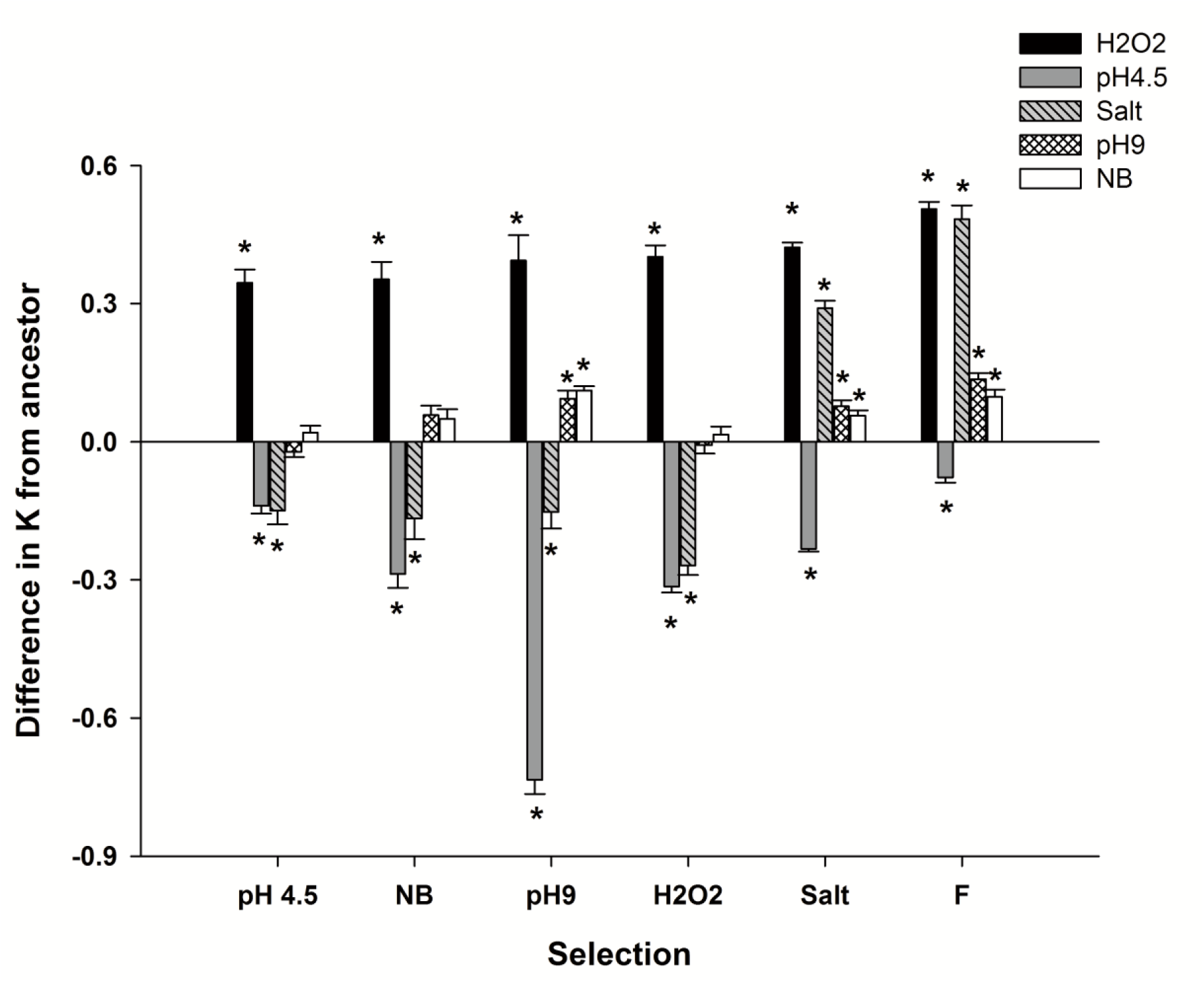
Difference (±SE) in K from ancestor for all the selection regimes. Difference in mean fitness (estimated as maximum density reached i.e. K) from ancestor was computed for every selection regime in every selection environment. Negative values indicate loss of fitness while positive values indicate gain of fitness, as compared to the ancestor. Every selection regime, except F, shows loss of fitness in at least one of the environments. * denotes *p* < 0.05 after individual ANOVAs followed by Holm-Šidák correction.

## DISCUSSIONS

### Complex, unpredictable fluctuations select for higher overall mean fitness

When subjected to predictable oscillations in a single environmental parameter over long time scales, microbial populations typically evolve to have higher fitness over the entire range of environments faced (Leroi et al. 1994; Turner and Elena 2000; Hughes et al. 2007; Coffey and Vignuzzi 2011; Alto et al. 2013; Puentes-Téllez et al. 2013; Condon et al. 2014). However, the evolutionary outcomes become much more diverse when the environment undergoes unpredictable fluctuations and the fitness of the selected populations can show no change (Alto et al. 2013), increase across the board (Turner and Elena 2000; Cooper and Lenski 2010; Ketola et al. 2013) or an increase with respect to a few life history traits with no change (Hughes et al. 2007) or decrease with respect to others (Hallsson and Björklund 2012). To further complicate matters, natural environments typically consist of multiple parameters that can change simultaneously (Lindow and Brandl 2003; Okafor 2011). Our results show that when subjected to such complex, unpredictable fluctuations for ~900 generations, the bacterial populations show modest in increase fitness in the stresses under which they evolved (Fig 1A).

Increasing the overall fitness across different environmental parameters is more challenging than to improve fitness for different values of the same environmental parameter. If successive environments are uncorrelated then there is little chance of a mutation beneficial in one environment to increase fitness in another. When mutations advantageous in a given selection environment have antagonistic pleiotropic effects in other selection environments (Cooper and Lenski 2000; Zhong et al. 2004; Duffy et al. 2006), they are negatively selected when the environment changes. Finally, in the absence of pleiotropic effects, only those mutations which are beneficial in some environments and neutral in other environments (i.e. conditionally neutral) can persist in a population over long time. Due to these constraints on the fixation of beneficial mutations in unpredictably fluctuating complex environments, it can be difficult for populations to show improvement in fitness for any given component of the environment. In line with these expectations, a previous study had reported little improvement in overall mean fitness when *E. coli* were selected in complex, unpredictable fluctuations over a short duration of ~170 generations (Karve et al. 2015). However, our results show that over a longer duration of selection, populations facing complex, unpredictable fluctuations can improve fitness over all the selection environments.

Apart from delaying the fixation of conditionally beneficial mutations, complex, unpredictable fluctuations are expected to strongly select against all those mutations that can reduce fitness in any of the selection environments (i.e. pH, salt or H_2_O_2_). This should reduce the loss of fitness across selection environments which, in turn, should lead to reduction in the variation for fitness. Our results for variation in fitness across the five selection environments support this expectation (Fig 1B).

### Fluctuating environments minimize the variation for fitness

When evolved under predictable or unpredictable temporal fluctuations, microbial populations show reduced variation for fitness over the whole range of selection environments (Kassen 2014 Table 4.2). This is because, when the environment changes temporally, it is the geometric mean (and not the arithmetic mean) of the fitness over the entire evolutionary time that plays a more important role in determining the long-term evolutionary success (Gillespie 1977). Since reducing the variation in a series increases its geometric mean (Orr 2007), populations facing fluctuating environments are expected to have lower variation in fitness across the selection environments. This prediction is supported by empirical studies where pH or host type fluctuates across time (Kassen 2014 Table 4.2). However, reducing the variation in fitness when multiple selection environments are changing unpredictably is expected to be much more challenging particularly when there are negative pleiotropic interactions between the traits. Interestingly, our results show that this is not the case as populations selected under complex, unpredictable fluctuations minimized the variation in the fitness across all selection environments (Fig 2).

The significantly lower variation for fitness in F populations compared to the ancestor needs to be interpreted together with the observation that the F populations also show significantly higher overall mean fitness (Fig 1A). This result indicates that F populations not only improved the fitness in some environments but also gained fitness in at least few selection environments. To confirm this, we estimated the differences in fitness under different environments from the ancestors.

### Fluctuating environment minimizes trade-off

All else being equal, when an ancestral population is close to the fitness maxima in a given environment, there will be little response to selection. On the other hand when the ancestor is far away from the fitness maxima for a given environment, given the required genetic variation, improvement in fitness in the selection environment is expected, even if it is accompanied by loss of fitness in other environments. However, improvement in fitness can be constrained in case of environmental heterogeneity (temporally fluctuating or complex or both), where populations face multiple environments during selection. If an increase in fitness in one component environment is accompanied by a decrease in fitness in another component environment, then the presence of such trade-off structures can potentially reduce the rate of adaptation, cause no adaptation or might even lead to maladaptation for some component environments. Thus, bacterial populations facing multiple environments (simultaneously or sequentially) are expected to show lesser increase in fitness as compared to that of the specialists.

However, results of empirical studies, involving both predictable and unpredictable fluctuations in single environmental parameter or host, do not agree with these predictions. Populations facing multiple values of a given environmental parameter can improve fitness over all or some of the selection environments without a loss of fitness in other selection environments (Turner and Elena 2000; Hughes et al. 2007; Condon et al. 2014). Our results extend this understanding to complex environments where multiple parameters fluctuate unpredictably over time. The populations selected under constant exposure to hydrogen peroxide and pH9 showed increase in fitness in the respective selection environments (Fig 2) which suggest that the ancestor was away from the fitness maxima of these two selection environments. However, this increase was accompanied by a loss of fitness in environments not experienced during selection (Fig 2). On the other hand, although the F populations showed increase in fitness for pH 9 and hydrogen peroxide, they did not lose fitness in any of the selection environments (Fig 2). In other words, the F populations not only adapted to these environments but could also by-pass the trade-offs that were experienced by the pH9 or hydrogen peroxide selected populations.

In case of pH4.5, neither F populations nor the populations selected under constant pH4.5 showed improved fitness over the ancestor (Fig 2). Thus, the ancestor was most likely well adapted to pH4.5 which is not surprising given that *Escherichia coli* are known to be well-adapted to acidic environments (reviewed in Foster 2004). Similarly, control environment of NB^Kan^ shows no improvement in fitness, which is intuitive given the ancestral *Escherichia coli* strain K12 is expected to be well adapted to the laboratory conditions.

The populations facing complex, fluctuating environments seem to face negligible constraints while adapting to component selection environments. The extent of adaptation seems to be governed by the distance of the ancestor from fitness maxima rather than the constraints imposed by the environment. In fact in salt, F populations show gain of fitness as compared to the ancestor while populations selected in constant exposure to salt display loss of fitness (Fig 2).

### Trade-off can be observed across traits for a given environment

This counterintuitive observation was resolved when we looked at a different proxy of fitness, namely, maximum density achieved or *K* (Novak et al. 2006). Populations selected in salt showed increased *K* relative to the ancestor (Fig 3). This improvement in *K,* and lack of the same in maximum growth rate as a result of selection, could be due to an underlying trade-off between growth-rates and carrying capacity. Higher concentration of salt will result in strong selection for robust membrane structures, which can be negatively correlated with the growth rate of the cells (Carlquist et al. 2012). Consistent with the observations for maximum growth rate as a proxy of fitness, even the F populations showed corresponding improvement in K. In the light of these results though, the choice of fitness proxy demands further attention.

In many studies on microbial experimental evolution, competitive ability relative to the ancestor is used as a proxy of fitness (Leroi et al. 1994; Hughes et al. 2007; Alto et al. 2013). This measure is preferred because it integrates over all phases of a growth cycle (i.e. lag phase, log phase etc.) and is expected to show how much better the evolved population has become in terms of evolutionarily replacing the ancestor (Kassen 2014 Page 16). However, we refrained from using this measure in the present study. This is because change in competitive fitness compared to the ancestor is an appropriate measure in case of constant or directionally changing selection environments, where populations are adapting towards a fixed fitness peak (Collins 2011). As opposed to this, adaptation to unpredictable environments will involve sudden changes in the underlying fitness landscape. Populations facing such environments will not show monotonic change in the relative fitness as compared to the ancestors. Thus, estimating the competitive fitness against the ancestor at any given point in the evolutionary trajectory does not give us much useful information about how much the population has really changed (Collins 2011). Incidentally, the same argument holds for any other measurement of fitness including the ones that we have used in this study (growth rate and carrying capacity). Therefore, these two indices are perhaps better viewed as fitness-related properties of the populations and in that sense are in the same epistemological category as competitive ability vis-a-vis the ancestors. Focussing on these two fitness-properties allows us to draw some interesting conclusions. The results of maximum growth rate and *K* are comparable in case of overall mean fitness and variation for fitness (Fig 1 and S4). But this is not the case when we consider the change in fitness from ancestor (Fig 2 and 3). Populations selected in salt show reduced maximum growth rate as compared to the ancestor but significantly higher K. Similarly in pH 4.5, maximum growth rate does not evolve in comparison to ancestor in all the selected populations but *K* shows a significant reduction (Fig 2 and 3). This shows that evolutionary interpretations based on different fitness properties may or may not agree with each other and different selection environments can select for different components of fitness. This is analogous to the concept of stability properties in ecology (Grimm and Wissel 1997) where it has been shown that evolution of one kind of stability may or may not lead to the evolution of another type (Dey et al. 2008). Thus, concentrating on any one fitness property in experimental evolution studies might lead to an under-estimation of the richness of the evolutionary process.

## Conclusions

To our knowledge, this is the first study which shows that populations can improve the fitness and minimize the variation for fitness when exposed to complex, unpredictable fluctuating environments. Simultaneous exposure can result in fitness increase in some selection environments without any loss of fitness in other selection environments. More importantly, our results suggest a possible explanation for the absence of trade-offs across environments in microbial populations by showing the other kind of trade-offs can exist i.e. trade-offs between different measures of fitness in the same environment. Estimating different components of fitness separately might be fruitful for future experimental studies.

## Acknowledgements

We thank Dr. Manjula Reddy for providing Kanamycin resistant strain of *Escherichia coli* K12. We thank Somendra Singh Kharola and S. Selweshwari for their help in laboratory work. SK was supported by a Senior Research Fellowship from Council of Scientific and Industrial Research, Govt. of India. This project was supported by a grant from Department of Biotechnology, Government of India and internal funding from Indian Institute of Science Education and Research, Pune. The authors declare no conflict of interests.

## Supplementary online material

**S1.Details of the ancestral Escherichia coli population used for the selection**

We used an *Escherichia coli* K12 MG1655 strain in which the lacY gene had been replaced with a Kanamycin resistance gene. Colonies of this bacterium are white coloured on MacConkey’s agar as opposed to the red coloured colonies produced by other *Escherichia coli*.

**S2.**
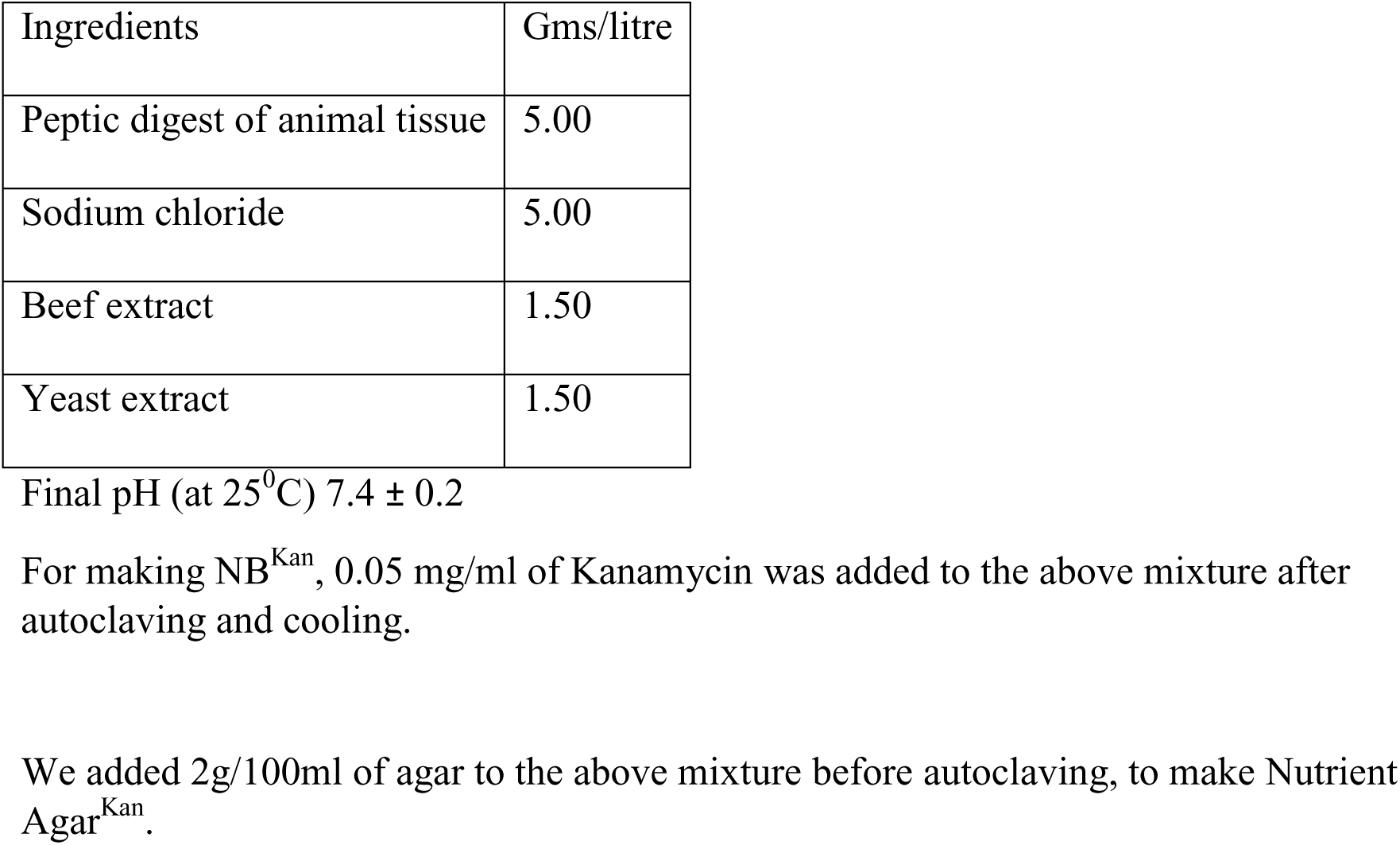
Composition of Nutrient broth -.

**S3.**
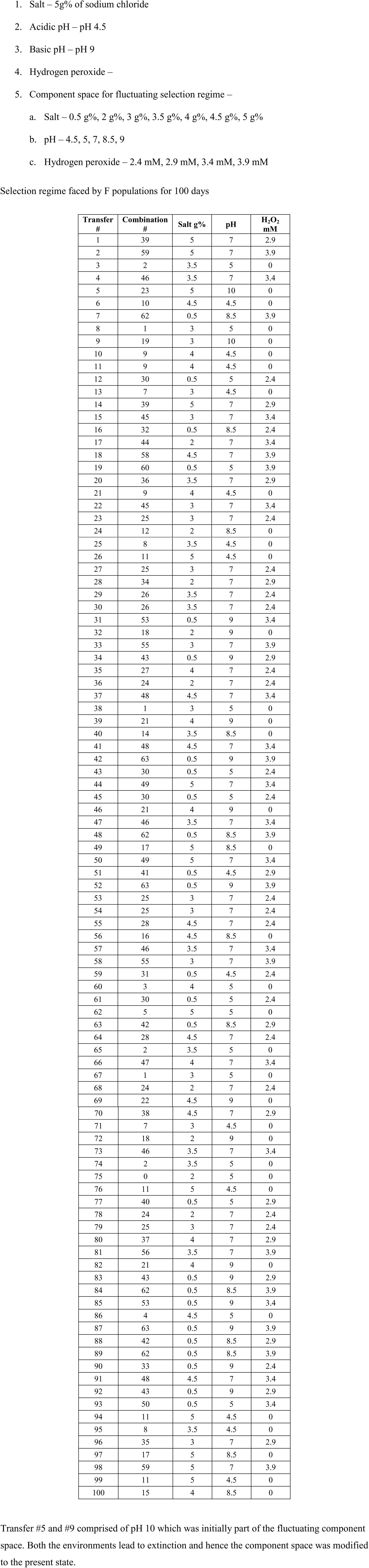
Details of all the selection regimes.

**S4:**
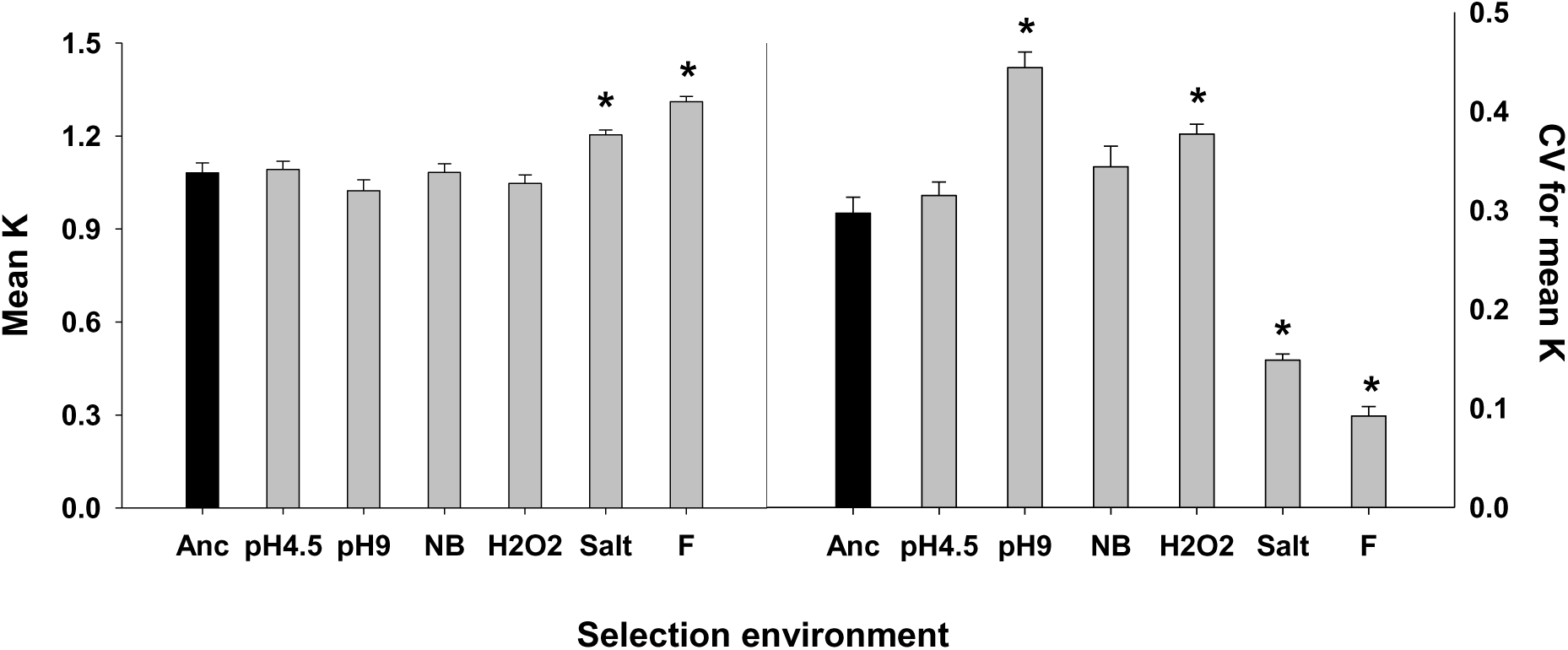
Mean *K* and coefficient of variation for *K*

**S5.**
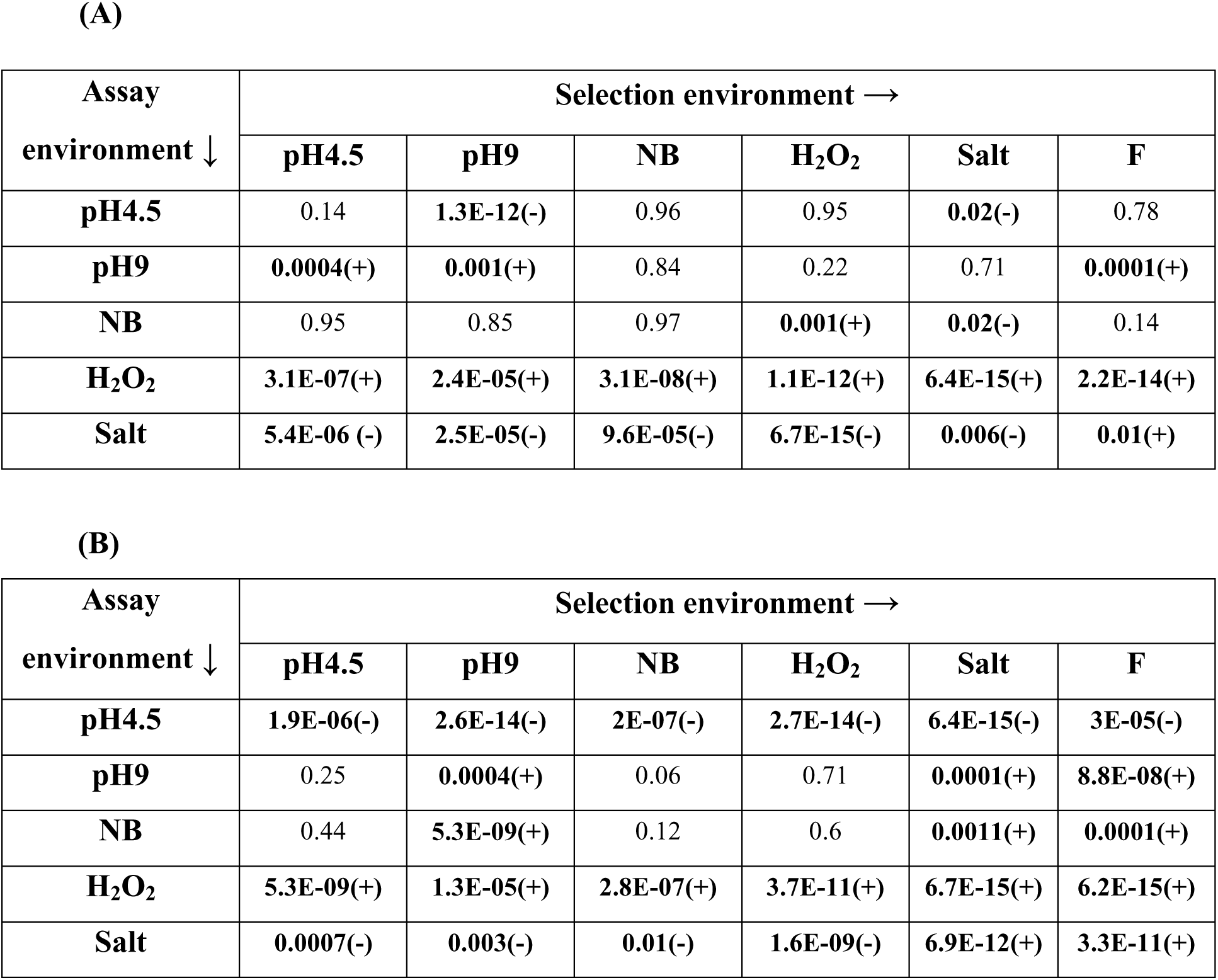
Difference from ancestor was computed for all the selection regimes in all the assay environments. Holm – Šidàk corrected *p* values for 30 t-tests are given for (A) maximum growth rate and (B) maximum density achieved. *p* < 0.05 denotes significant difference from ancestor which is accompanied by a sign in the bracket denoting direction of change. ‘-’ represents decrease while ‘+’ represents increase in fitness from ancestor.

